# Enhanced pitch perception in early blind individuals and musicians is due to reduced internal noise

**DOI:** 10.1101/2025.11.25.690447

**Authors:** Woon Ju Park, Jasmine Awad, Ione Fine

## Abstract

Although it is commonly assumed that early blind individuals have enhanced pitch perception, the literature to date has been contradictory. We measured performance in two tasks known to recruit early vs. late stages of auditory frequency tuning in early blind and sighted individuals with matched levels of musical training. The two groups showed equivalent performance on a tone detection in a notched-noise task that assesses peripheral and subcortical frequency selectivity. In contrast, early blind individuals showed enhanced performance in a pitch discrimination task thought to be mediated by cortical mechanisms. Computational modeling revealed that this enhancement was best explained by reduced internal noise rather than narrower frequency tuning, with both blindness and musical training predicting a reduction in internal noise. These findings identify internal noise as a key factor in experience-driven auditory plasticity as a result of both musical experience and early blindness.

## Introduction

For centuries, it has been postulated that individuals who become blind early in life develop superior pitch discrimination abilities. Indeed, a remarkable number of blind individuals have careers as piano tuners and musicians. However, while this enhanced pitch perception is commonly cited as a classic example of experience-dependent auditory plasticity, the empirical evidence supporting this phenomenon remains mixed (Sabourin et al., 2022). Some studies report enhanced performance (Gougoux et al., 2004; Voss & Zatorre, 2012; Wan et al., 2010), whereas others find performance comparable to sighted controls (Collignon et al., 2013; Watkins et al., 2013).

Pitch perception relies on multiple stages of auditory processing. At the periphery and brainstem, cochlear filters determine the precision with which different frequencies are resolved (Glasberg & Moore, 1990; Patterson, 1976; Shera et al., 2002). Even at these early stages, frequency selectivity can be shaped by top-down effects. Corticofugal feedback can modulate cochlear responses (Terreros & Delano, 2015), and microstimulation of the auditory cortex alters the otoacoustic emissions from the auditory periphery in humans (Perrot et al., 2006). Experience-dependent plasticity has also been implicated here: professional musicians—another population with extensive auditory experience—show sharper cochlear frequency tuning, with tuning strength correlated with years of musical training (Bidelman et al., 2014, 2016).

Beyond these early stages, cortical circuits integrate spectral and temporal information to construct the percept of pitch (Bendor & Wang, 2005; Oxenham, 2012). Enhancement at this stage could arise through several mechanisms. One possibility is sharpened frequency tuning. Early experience shapes cortical frequency selectivity in animals, with the elimination of patterned acoustic input extending the critical period (Chang et al., 2005; Chang & Merzenich, 2003), and extensive exposure to a single frequency resulting in an expansion of the representation of that frequency (De Villers-Sidani et al., 2007; Zhang et al., 2001). In line with this, in early blind individuals, a very small narrowing of auditory frequency tuning has been observed in the primary and secondary auditory cortices using fMRI-based population receptive field mapping (Huber et al., 2019).

Another way auditory processing can be enhanced at the cortical stage is via reduced internal noise. Internal noise manifests as trial-by-trial variability in neural responses (Arazi et al., 2017; Carandini, 2004; Churchland et al., 2010). Lower internal noise would allow more consistent encoding of pitch differences, improving discrimination without changes in frequency tuning per se. Indeed, studies on training-induced perceptual learning have shown that discrimination abilities improve not through sharpening of receptive fields but through enhancement of the readout of relevant neural populations: increasing the weighting of informative neural signals and suppressing task-irrelevant activity, which effectively adds noise, enhances perceptual performance (Bejjanki et al., 2011; Dosher & Lu, 1998).

To address how and where auditory enhancements might arise in early blind individuals, we measured performance on two psychoacoustic tasks that probe frequency selectivity at different stages of the auditory system. Our goal was to test which level of processing—peripheral/brainstem vs. cortical—shows evidence of enhancement, and through what mechanism such enhancement might occur.

The first was tone detection in notched noise (Figure 1A). Participants indicated which of the two intervals contained a signal tone (1200 Hz) embedded in broadband noise (300 – 4800 Hz). The noise had a spectral notch around the signal frequency, whose width was varied to adjust task difficulty. Threshold and attentional lapse rate were measured at each notch width. Because performance depends on how sharply cochlear filters separate frequencies, the resulting “behavioral tuning curve” is interpreted as a proxy for early auditory filter bandwidth (Glasberg & Moore, 1990; Patterson, 1976).

**Figure 1.**
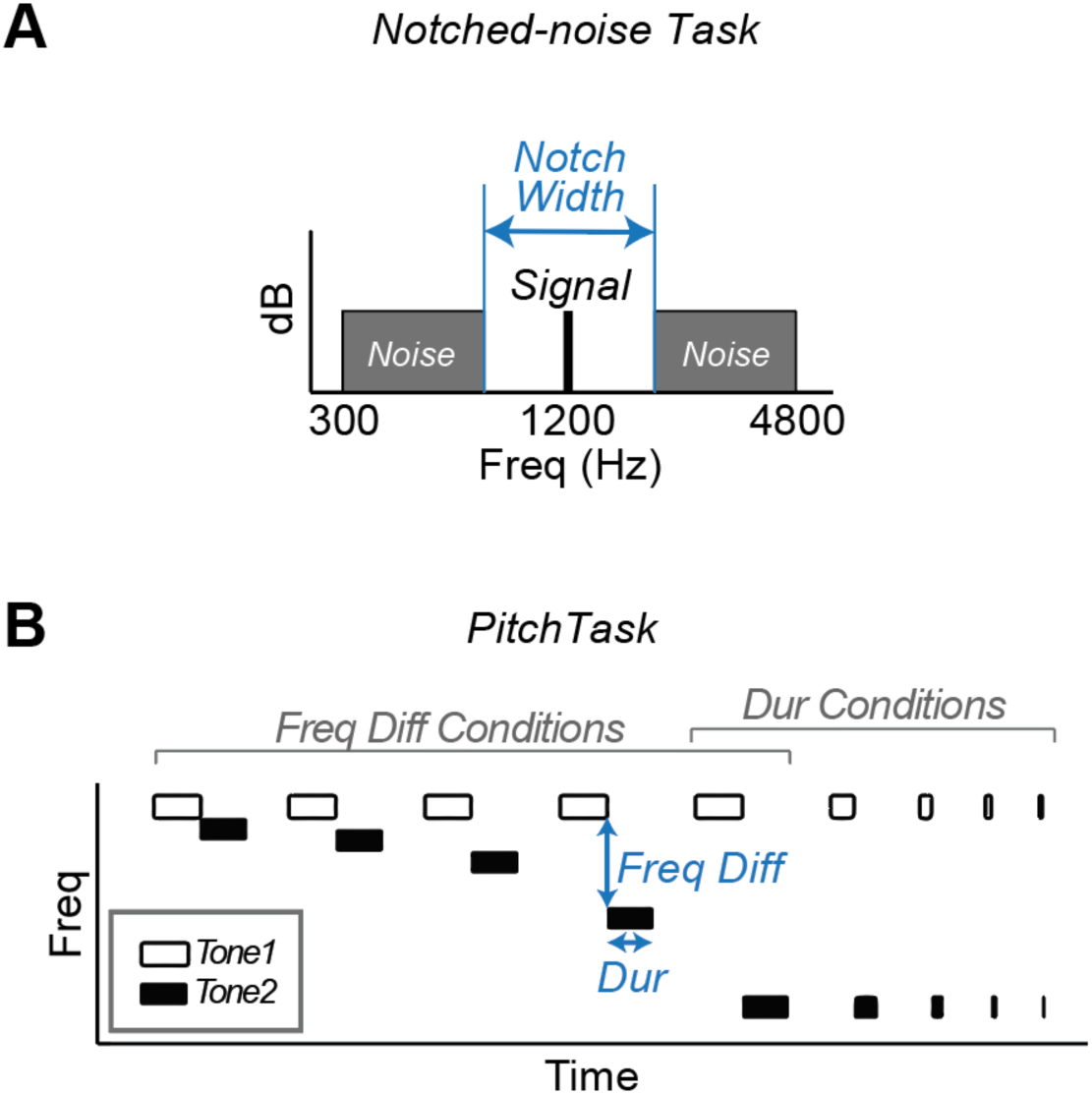
Methods. A. Notched-noise task. Participants were asked to detect a signal tone (1200 Hz) embedded in two notched noise bands (300 - 4800 Hz). The notch width (frequency difference between the two noise bands) was manipulated to adjust difficulty. Participants indicated which interval contained the signal tone. B. Pitch task. Participants discriminated between two consecutively presented pure tones and reported whether the two tones were rising (the second sound higher) or falling (the second sound lower) in pitch. The frequency difference between the two tones or the duration of the tones was varied to adjust task difficulty.

Our second task, pitch discrimination (Figure 1B), is likely mediated by the primary auditory cortex (Abrams et al., 2025; Bendor & Wang, 2005, 2010). Participants judged whether the pitch of two consecutively presented tones was rising or falling. Following Gougoux et al. (Gougoux et al., 2004), who found some of the earliest behavioral evidence for enhanced pitch perception in early blind individuals, task difficulty was adjusted by varying both the frequency difference between the two tones (at a fixed duration) as well as the tone duration (at a fixed frequency difference). Accuracy was measured for each of the durations and frequency differences.

We tested 9 early blind and 18 sighted participants. To control for musical experience, two sighted participants were matched to each early blind participant based on their age and years of musical training (see Table 1 for the early blind participant characteristics). Importantly, by equating musical experience between the groups (sighted and early blind), the design allowed us to attribute any group differences to early blindness itself, while also making it possible to assess how musical training contributes to auditory performance independently of blindness.

**Table 1.** Early blind participant characteristics.

| ID | Sex | Age | Blindness<br>Onset | Cause of Blindness | Light<br>Perception | Years Musical<br>Training |
| --- | --- | --- | --- | --- | --- | --- |
| EB1 | M | 37 | Born blind | Leber's congenital<br>amaurosis | Low | 0 |
| EB2 | F | 60 | Born blind | Retinopathy of<br>prematurity | No | 10 |
| EB3 | M | 56 | Born blind | Congenital glaucoma | No | 50 |
| EB4 | M | 29 | Age 2 | Retinoblastoma | No | 22 |
| EB5 | F | 58 | Age 2 | Optic atrophy | No | 50 |
| EB6 | F | 32 | 8 Months | Retinoblastoma | No | 10 |
| EB7 | F | 43 | Born blind | Congenital glaucoma | No | 4 |
| EB8 | M | 67 | Born blind | Retinopathy of<br>prematurity | No | 59 |
| EB9 | M | 49 | Born blind | Leber's congenital<br>amaurosis | Low | 41 |

Our findings show a clear dissociation in the two tasks: Early blind and sighted individuals performed comparably on the notched-noise task, providing no evidence for enhanced frequency selectivity at peripheral or brainstem levels. In contrast, early blind individuals showed significantly better performance on the pitch discrimination task. Computational modeling of performance in the pitch task suggests that the enhanced performance found for both early blind individuals and individuals with many years of musical training was mediated by reduced internal noise rather than narrower frequency tuning. Notably, the effect of early blindness on internal noise was not explained by musical experience alone, demonstrating that blindness contributes uniquely and independently to reduced internal noise in pitch processing at higher levels of the auditory pathway.

## Results

### No significant difference between early blind and sighted individuals for the notched-noise task

The early blind and sighted individuals showed similar performance for detecting a signal tone embedded in notched noise (Figure 2A). The effects of blindness and notch width on thresholds were assessed using a mixed-effects linear model analysis with group, notch width, and their interaction as fixed effects. The analysis also accounts for the individual matching between early blind and sighted participants by including the subject-level pairing between the two groups as a random intercept (see Methods). The group difference in attentional lapse rates was tested using a non-parametric Wilcox rank-sum test.

**Figure 2.**
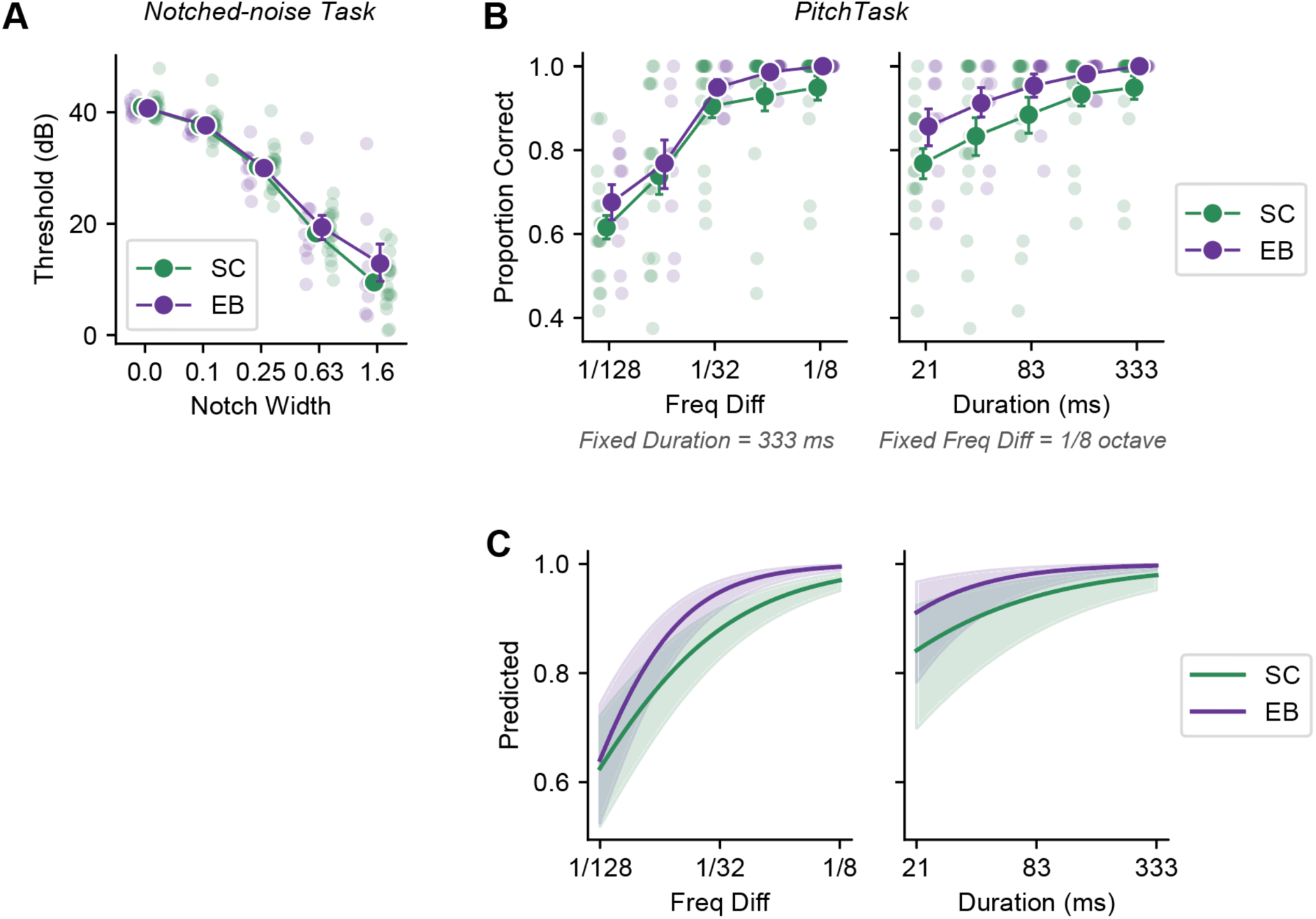
Task performance. A. Notch task. Early blind (EB) individuals and sighted controls (SC) showed comparable performance in the notch task, demonstrating overall increasing sensitivity (lower thresholds) as the notch width became larger. B. Pitch task. Both EB and SC showed decreasing performance as the task became more difficult, but EB individuals performed better than SC for easier conditions when the frequency difference was larger (left panel) and stimulus duration was longer (right panel). Colored circles show group means. Error bars are standard errors of the mean. C. Predicted performance from the mixed effects logistic regression models. Shaded regions represent 68% confidence interval.

As expected, both early blind and sighted groups showed increasing sensitivity (i.e., lower thresholds) as the notch width became wider (F(4, 100) = 320.87, p < .001). However, the two groups had comparable sensitivity (F(1, 17) = .36, p = .56), and there was no interaction between group and notch width (F(4, 100) = 1.23, p = .30). The two groups had similar attentional lapse rates (EB: mean = 0.0036, SD = 0.0031; SC: mean = 0.0057, SD = 0.0039; W = 54, p = .18). These results suggest that there is little effect of early blindness on frequency tuning within early stages of auditory processing.

Given the previous reports of sharper cochlear tuning in professional musicians (Bidelman et al., 2014, 2016), we additionally examined whether musical training predicted performance in the notched-noise task. Unlike prior reports, years of musical training did not predict overall average threshold (EB: r(7) = −.18, p =.64; SC: r(16) = .11, p = .67) nor did it predict the slope in the tuning curve (the difference in thresholds between the smallest and largest notch widths; EB: r(7) = .11, p = .79; SC: r(16) = .16, p = .51) for either early blind or sighted individuals. Thus, neither early blindness nor musical training enhanced behavioral tuning at the early stages of auditory frequency selectivity.

### Early blind individuals can discriminate pitch better than sighted individuals

In contrast, early blind individuals were better than sighted controls at the pitch task. Because the dependent variable was the proportion of correct responses, we used mixed effects logistic regression models (see Methods) to separately explain performance as a function of frequency difference (for a fixed tone duration of 333 ms) and as a function of tone duration (for a fixed frequency difference of 1/8 of an octave) conditions. Figure 2B shows the observed data, and Figure 2C shows the logistic regression model fits.

As expected, for the frequency difference conditions (Figures 2B & 2C, left), performance for both groups declined as the frequency difference between the two tones became smaller (effect of frequency difference: χ^2^(1) = 1.20, p < .001). The main effect of group was not significant (χ^2^(1) = 1.20, p = .27). However, there was a significant interaction between frequency difference and group (χ^2^(1) = 9.64, p = .002). Early blind individuals performed better than sighted controls when the frequency difference was larger.

In the duration conditions (Figures 2B & 2C, right), as expected, performance for both groups declined as the duration of the sounds became shorter (effect of duration: χ^2^(1) = 131.32, p < .001). There was a trend toward a main effect of group, in which early blind individuals overall performed better than sighted controls: χ^2^(1) = 3.50, p = .061. Similar to the frequency difference conditions, we also observed a significant interaction between group and duration: χ^2^(1) = 5.09, p = .024. Specifically, the better performance of early blind individuals was more pronounced for longer stimulus durations. These findings suggest a clear dissociation, with early blindness enhancing pitch discrimination but not early stages of auditory processing.

### Enhanced pitch discrimination in early blind individuals can be explained by reduced internal noise

To determine through what mechanism—frequency tuning vs internal noise—pitch discrimination performance can be enhanced via early blindness at higher levels of auditory processing, we used a simple model based on signal detection theory (see Methods for details) to fit behavioral performance in the pitch task as a function of frequency and duration. The model assumes that the internal neural representation of each of the two tones is determined by two parameters that determine frequency tuning (σ_f_), and internal noise (σ_n_). On each trial, the listener compares the internal responses resulting from mechanisms tuned to the higher and lower frequencies, and responds based on which mechanism had the larger response. The internal response is generated by averaging a noisy signal over time, so longer stimulus durations lead to less noisy responses and better performance.

We fit this signal detection model to each participant’s pitch discrimination data (both frequency difference and duration conditions combined), with two free parameters representing frequency tuning width (σ_f_) and internal noise (σ_n_). Representative model fits for an early blind and sighted participant are shown in Figure 3A. The model successfully predicted pitch discrimination performance as a function of frequency difference and duration across individuals. The mean correlation coefficient between observed and predicted performances was 0.96 for the early blind (SD = 0.04), and 0.87 for the sighted (SD = 0.21) groups, with no evidence of any systematic deviation between model predictions and data, Figure 3B.

**Figure 3.**
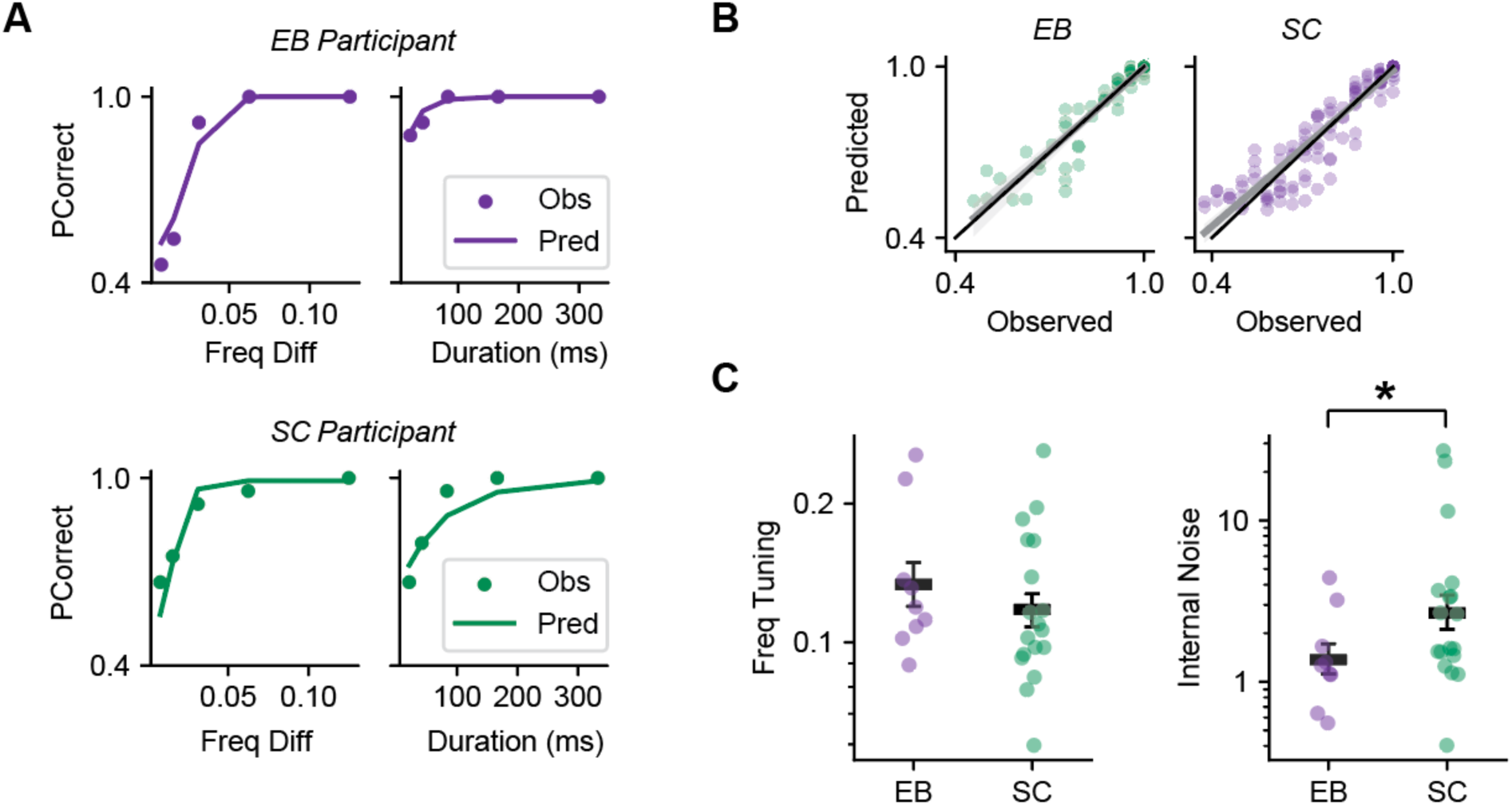
Understanding the variables influencing pitch performance. A. Example of model predictions for a representative EB (top) and SC (bottom) participant. Colored circles show observed data, and the lines show model predictions. B. Fitted model predictions in comparison to the observed performance. Each dot represents each condition for an individual participant. Black and gray lines are the identity line and regression fits across all data points, respectively. C. Parameter estimates. The two groups showed no difference in the frequency tuning (σ_f_; left). In contrast, EB individuals had significantly reduced internal noise (σ_n_) compared to SC (right). Colored circles show individual data and black horizontal bars show group means. Error bars are standard errors of the mean.

Best-fitting parameter estimates (Figure 3C) showed no significant difference in frequency tuning between early blind and sighted groups (t(17) = 0.98, p = 0.34). In contrast, the parameter representing internal noise was significantly lower in early blind individuals as compared to sighted controls (t(17) = −3.20, p = 0.005). To formally test how much either parameter contributed to explaining group differences, we compared the full model, in which both σ_f_ and σ_n_ varied across individuals, to reduced models, in which either σ_f_ or σ_n_ was fixed to a single value (shared across groups) while the other was allowed to vary across individuals. Likelihood ratio tests showed that fixing frequency tuning resulted in a modest loss of fit relative to the full model (Δ–2LL (full - reduced) = 10.62, χ²(1) = 10.62, p = .0011; ΔAIC (full - reduced) = −8.62). By contrast, fixing internal noise led to a much larger deterioration in fit (Δ–2LL = 68.33, χ²(1) = 68.33, p < .001; ΔAIC = −66.33). These model comparisons indicate that reduced internal noise, rather than altered frequency tuning, provides the primary explanation for the superior pitch discrimination observed in early blind participants.

### Reduced internal noise is associated with both blindness and longer years of musical training

The reduced internal noise observed in early blind individuals might be explained either by loss of vision or as a result of extensive auditory experience. To address this, we began by examining whether musical training was related to the signal detection model parameters. For frequency tuning parameter (Figure 4A), years of musical training did not correlate with tuning in either of the groups (early blind: r(7) = 0.60, p = 0.08; sighted: r(16) = −0.12, p = 0.64). In contrast, internal noise showed a significant negative correlation with musical training in both early blind (r(7) = −0.70, p = 0.03) and sighted groups (r(16) = −0.54, p = 0.02; Figure 4B). This suggests that individuals with longer years of musical training tend to have lower internal noise.

**Figure 4.**
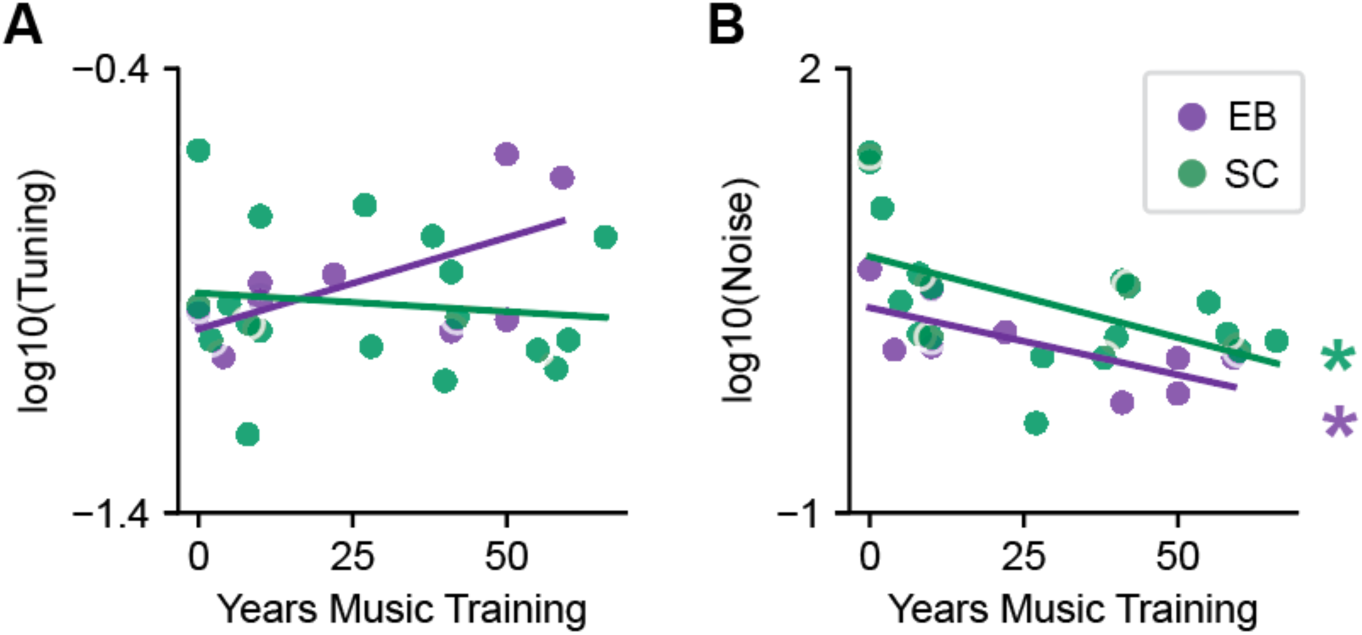
Effects of music training on parameter estimates. A. The estimated frequency tuning (σ_f_) was not significantly correlated with years of music training across individuals in either of the groups. B. However, there was a significant correlation between estimated internal noise (σ_n_) and years of music training in both groups, where longer music training was associated with lower internal noise.

However, as can be seen in Figure 4B, the effect of blindness cannot be explained by years of musical training alone. To test whether early blindness explained additional variance in internal noise beyond musical experience, we fit a mixed-effects linear regression model that included both years of musical training and group (early blind vs sighted) as fixed effects, with random intercepts for matched pairs.

The analysis again showed that internal noise decreased with longer years of musical training (β = −.011, SE = .004, t(10.27) = −2.76, p = .019). Critically, early blindness remained a significant predictor even after controlling for musical training (β = .29, SE = .096, t(17.88) = 3.05, p = .007), demonstrating that the effect of early blindness is not explained by differences in musical experience. When the interaction between group and musical training was included as a predictor in the model, the interaction effect was not significant (β = −.0014, SE = .004, t(18) = −.32, p = .75), and including an interaction term did not improve the model fit: a model without the interaction was preferred (AIC = 21.19) over the model with the interaction (AIC = 23.09; ΔAIC (full – reduced) = 1.9). A likelihood-ratio test comparing the nested models similarly found no improvement from adding the interaction (χ²(1) = 0.10, p = .75). Thus, both early blindness and musical experience independently reduce internal noise, and the reduced noise observed in early blind listeners cannot be explained solely by musical training.

## Discussion

Our findings demonstrate that early blindness is associated with enhanced pitch perception. This enhancement likely arises from higher stages of auditory processing, as no group differences were observed in a task designed to assess peripheral and subcortical frequency selectivity. Notably, a similar enhancement was observed in individuals with extensive musical training, suggesting that both early blindness and musical experience can lead to improved pitch discrimination. Computational modeling further revealed that superior pitch perception in both groups is best explained by reduced internal noise, rather than by narrower frequency tuning. This challenges the common assumption that auditory plasticity—whether driven by blindness or by musical expertise—primarily reflects sharpened frequency tuning in the auditory cortex.

### No effects of early blindness or musical training in frequency selectivity measured by the notched-noise task

We found no differences in frequency selectivity between early blind and sighted participants in the notched-noise task. Nor did we see an effect of musical training in either group. This finding was somewhat unexpected, given previous work showing that professional musicians exhibit sharper cochlear-level frequency selectivity, which is correlated with years of musical experience (Bidelman et al., 2016). However, reanalysis of the data in this earlier study suggests that the finding of a significant correlation between years of musical experience and cochlear tuning was primarily driven by the contrast between musicians and non-musicians, since all non-musicians had less than three years of musical training. The correlation with years of experience *within* the musician group seems to have been significant near the 0.05 level with 14 participants.

In our study, because our goal was to match musical training between early blind and sighted participants, we selectively recruited non-professional musicians who nonetheless had many years of musical practice/training. While this group was self-reported, we defined being in training as 1-2 hours of regular practice per week. Our failure to find an effect of musical experience suggests that the findings of Bidelman et al. may reflect a selection bias rather than an effect of experience. Being a professional musician is a challenging career which is often financially unsustainable. Those who reported still being a professional musician after 15-20 years are likely to have been particularly talented, which may have included sharper cochlear tuning.

Our finding that early blind individuals outperform sighted controls only in pitch discrimination, and not tone detection in notched noise, allows us to exclude several possible explanations for differences in pitch selectivity. First, our failure to find a difference in the notch task challenges the simplistic notion of across-the-board “enhanced hearing” in the absence of vision. On a related note, it excludes the possibility (always a concern in a small sample study) that we selectively recruited early blind individuals with particularly good hearing. Finally, it excludes the possibility that group differences were due to high-level cognitive differences – for example, early blind individuals might have been more capable of prolonged attention to an auditory task.

### Enhanced pitch discrimination in early blind individuals depends on stimulus conditions

The enhanced performance of early blind individuals for pitch discrimination was only found for larger frequency differences or longer stimulus durations. This may help explain the apparent inconsistencies observed in previous studies. Our task was based on an earlier study (Gougoux et al., 2004) that reported enhanced pitch discrimination in early blind individuals, although this study did not control for musical training. A later study, also based on the methods of Gougoux et al., examining pitch perception in a small number of individuals with anophthalmia failed to replicate this finding, and similarly did not account for musical training (Watkins et al., 2013). In contrast, studies that matched musical training across groups found a pattern consistent with ours: enhanced performance in early blind people for larger frequency differences (0.5 – 2%; (Wan et al., 2010)) and longer stimuli (300 ms; (Voss & Zatorre, 2012)), but not for shorter durations (150 ms; (Collignon et al., 2013)).

### Enhanced pitch perception in both early blind individuals and musicians seems to be mediated by reduced internal noise

Unexpectedly, enhanced pitch discrimination in early blind individuals was best explained by reduced internal noise, and not by sharpened frequency tuning. Although a previous study did find reduced tuning widths in the primary and secondary auditory cortices of early blind individuals using fMRI (Huber et al., 2019), these reductions were very small – on the order of 3%. Thus, while there may be small reductions in frequency tuning bandwidth as a result of early blindness, any such reductions in tuning seem to not be the primary cause of improved pitch perception. Instead, our modeling points to reduced trial-by-trial variability in internal responses as the key mechanism. Internal noise, thought to reflect inherent fluctuation in neural activity (Carandini, 2004; Churchland et al., 2010; Dinstein et al., 2015; Goris et al., 2014; Shadlen & Newsome, 1994; Tomko & Crapper, 1974) constrains perceptual reliability (Arazi et al., 2017). Lower internal noise can improve the precision of auditory representations and thus discrimination performance.

This interpretation aligns with evidence from perceptual learning, which shows that training can enhance discrimination by improving the readout of task-relevant neural populations and down-weighting irrelevant ones, effectively reducing internal noise, while leaving the tuning of receptive field properties unaltered (Bejjanki et al., 2011; Dosher & Lu, 1998; Law & Gold, 2008). Our findings suggest that a similar enhancement may occur as a result of both musical training and early blindness. The idea that blindness enhances auditory perception via reduced internal noise is also consistent with recent work showing improved auditory motion perception in noise, where early blind individuals outperformed sighted controls by relying on spatiotemporal filters more precisely matched to the stimulus (Park & Fine, 2023).

In conclusion, our findings show that enhanced pitch discrimination in both early blind individuals and musicians is well explained by reduced internal noise that occurs at later stages of auditory processing. In blind individuals this reduction of internal noise is likely to influence performance across a wide variety of auditory tasks. One limitation of our study is that it is impossible to determine whether the effect of blindness is due to the loss of vision or the increased reliance on auditory cues that occurs as a result of the loss of vision. One way to separate these might be to examine whether perceptual enhancements as a result of early blindness are more pronounced for stimuli that are ecologically critical for early blind individuals.

## Methods

This study was approved by the University of Washington’s Institutional Review Board and conducted in accordance with the Code of Ethics of the Declaration of Helsinki. Informed written consent was obtained from all participants prior to conducting the experiments.

### Participants

Participants included 9 early blind individuals (Table 1) and 18 sighted controls with neurotypical auditory histories. Two sighted individuals were matched to each early blind participant based on their age (within 10 years) and music experience (determined based on the self-reported number of years in musical training).

### General Methods

Auditory stimuli were delivered through Etymotic ER-2 insert earphones at a sampling rate of 44,100 Hz. The stimuli were generated and presented using MATLAB and Psychophysics Toolbox.

### Notch Task

#### Stimulus

The stimuli consisted of a target signal and a noise mask. The signal was a pure tone presented at 120 Hz for 300 ms. The noise mask was created by band-pass filtering broadband noise (300 – 4800 Hz, 40 dB) in the frequency domain. The ‘notch’ in this noise was centered on the signal tone, and the notch width ranged from 0 to 1.6 times the signal frequency. The noise mask was presented for 600 ms and temporally overlapped the stimulus (which was presented 150ms after mask onset). Both the signal and noise mask had 100 ms linear ramps at onset and offset and were presented binaurally.

A two-interval forced-choice procedure with adaptive signal-amplitude adjustment was used to determine the threshold for detecting the signal within the notched-noise. The two intervals, both containing the noise mask, were presented consecutively, separated by 300 ms of silence. One randomly chosen interval also contained the signal. The amplitude of the signal was adaptively adjusted using two interleaved one-up three-down staircases. Each staircase contained 48 trials, such that a session consisted of 96 trials total for each notch width.

#### Task

On each trial, participants reported which interval contained the signal by pressing the corresponding button on a keyboard. Auditory feedback (a brief beep) was provided if they were correct. Participants first completed 80 trials of practice and then two sessions, each consisting of 480 trials.

#### Analysis

Each participant’s threshold to detect the signal tone, as well as the lapse rate at each notch width, was estimated by fitting a Weibull function using the psignifit toolbox (Schütt et al., 2016), using all trials from each staircase within a session. This resulted in 4 threshold measurements, which were then averaged. A linear mixed-effect model analysis was performed to predict thresholds with notch width, group, and their interactions as fixed effects. The early blind and sighted pairing was included as random intercepts, allowing us to account for individual matching between early blind and sighted groups. The significance of the fixed effects was estimated via F tests using the Satterthwaite method. The effect of early blindness on attentional lapse rate was assessed using a Wilcox rank-sum test. Pearson correlation analyses were used to test the relationship between years of music experience on the overall performance (average threshold) and the slope (difference between the threshold at the smallest and largest notch widths) separately in each group.

### Pitch Task

#### Stimulus and Task

Participants listened to tone pairs binaurally at comfortable loudness levels and indicated via a button press whether the tone pair was rising (pitch in the second tone higher than the first) or falling (pitch in the second tone lower than the first) in pitch. At each of the four frequency levels (reference frequency: range, 500 – 1200 Hz), task difficulty (9 levels) was manipulated by parametrically varying either the frequency difference between the two tones (1/8, 1/16, 1/32, 1/64 and 1/128 of an octave) or the tone durations (333, 167, 83, 42, and 21 ms). There were 72 trials total within a block (4 frequency levels x 2 pitch directions x 9 difficulty levels), and participants completed three blocks.

#### Analysis

The percentage correct score was calculated across all levels of difficulty. Frequency difference and duration conditions were analyzed separately. In both analyses, mixed effects logistic regression was performed to predict the percentage correct scores with the log of either frequency difference or duration, group, and their interactions as fixed effects, and a combination of pair and subject as random intercepts (to account for individual matching between groups). Wald chi-square tests were then performed to understand the significance of fixed effects.

#### Signal Detection Model

To predict each participant’s response on each trial, we used a signal detection theory-based model that estimated the width of each participant’s frequency tuning and the level of internal noise. On each trial, the participant’s internal estimates of presented tones were derived from Gaussian tuning curves centered on the reference and the test frequency, with the standard deviation (σ_f_) as a free parameter determining the tuning width. Internal noise, randomly drawn from a zero-mean Gaussian distribution was then added to the estimates. The standard deviation of this noise distribution (σ_n_) was a free parameter that determined the level of internal noise. The final level of internal noise was determined by averaging noisy sample estimates over time. Thus, the longer the stimulus duration, the larger the number of samples that were averaged, resulting in a lower final level of internal noise.

The estimates for the two tones were compared against each other to predict the participant’s response. Correct responses occurred when the internal response for the reference frequency was larger than the response to the test frequency (we assume in this model that the order of intervals was unimportant). The procedure was repeated 10,000 times to derive the predicted proportion correct for each frequency difference and stimulus duration combination. When fitting the data for individual participants the two free parameters describing the standard deviations for the tuning curve and the Gaussian noise distributions, respectively, were estimated as values that maximized the likelihood of the participant’s responses.

To assess whether group differences in pitch discrimination were better explained by frequency tuning or internal noise, we fit signal detection theory–based models to data from both groups. The full model included separate parameters for frequency tuning (σ_t_) and internal noise (σ_n_) for early blind and sighted groups (4 free parameters). We then constructed two reduced (nested) models (3 free parameters each): one in which frequency tuning was fixed to a single parameter that was shared across groups while internal noise was free to vary across individuals, and another in which internal noise was fixed, while tuning was free to vary across individuals.

We compared each reduced model against the full model using likelihood ratio tests (LRTs). Specifically, we calculated the change in deviance (Δ–2LL) between models, which is χ²-distributed with degrees of freedom equal to the difference in the number of free parameters. We also computed Akaike information criterion (AIC) values to compare relative model fit while penalizing model complexity. In this framework, a significant LRT (p < .05) indicates that the additional free parameter in the full model provides a significantly better account of the data than the reduced model, whereas AIC differences > 10 are generally taken as strong evidence favoring the lower-AIC model.

### Statistical Analysis

For statistical analyses, we tested the effects of group and conditions at the significance level of .05. Holm–Bonferroni corrections were used for multiple comparisons as needed.

## Data Availability

The current IRB at the University of Washington restricts publicly sharing data that contain individual visual and medical histories. Data, in anonymized format, are available from the corresponding author upon request.

## Code Availability

Code for the experiments and analyses is openly available at https://github.com/VisCog/pitch-notch.

## Acknowledgements

Supported by the NIH R00-EY034546 to WJP, R01-EY014645 to IF, Weill Neurohub Postdoctoral Award to WJP, and Georgia Tech Smithgall-Watts Early Career Award to WJP.

## Author Contributions

Conceptualization: WJP, and IF; Methodology: WJP and IF; Software: WJP; Formal analysis: WJP; Investigation: JA and WJP; Writing—Original draft: WJP; Writing—Review & Editing: WJP, JA, and IF; Funding acquisition: WJP and IF.

## References

Abrams, E. B., Marantz, A., Krementsov, I., & Gwilliams, L. (2025). Dynamics of Pitch Perception in the Auditory Cortex. The Journal of Neuroscience, 45(12), e1111242025. 10.1523/JNEUROSCI.1111-24.2025

Arazi, A., Censor, N., & Dinstein, I. (2017). Neural Variability Quenching Predicts Individual Perceptual Abilities. The Journal of Neuroscience, 37(1), 97–109. 10.1523/JNEUROSCI.1671-16.2016

Bejjanki, V. R., Beck, J. M., Lu, Z.-L. L., & Pouget, A. (2011). Perceptual learning as improved probabilistic inference in early sensory areas. Nature Neuroscience, 14(5), 642–648. 10.1038/nn.2796

Bendor, D., & Wang, X. (2005). The neuronal representation of pitch in primate auditory cortex. Nature, 436(7054), 1161–1165. 10.1038/nature03867

Bendor, D., & Wang, X. (2010). Neural Coding of Periodicity in Marmoset Auditory Cortex. Journal of Neurophysiology, 103(4), 1809–1822. 10.1152/jn.00281.2009

Bidelman, G. M., Nelms, C., & Bhagat, S. P. (2016). Musical experience sharpens human cochlear tuning. Hearing Research, 335, 40–46. 10.1016/J.HEARES.2016.02.012

Bidelman, G. M., Schug, J. M., Jennings, S. G., & Bhagat, S. P. (2014). Psychophysical auditory filter estimates reveal sharper cochlear tuning in musicians. The Journal of the Acoustical Society of America, 136(1), EL33–EL39. 10.1121/1.4885484

Carandini, M. (2004). Amplification of Trial-to-Trial Response Variability by Neurons in Visual Cortex. PLoS Biology, 2(9), e264. 10.1371/journal.pbio.0020264

Chang, E. F., Bao, S., Imaizumi, K., Schreiner, C. E., & Merzenich, M. M. (2005). Development of spectral and temporal response selectivity in the auditory cortex. Proceedings of the National Academy of Sciences, 102(45), 16460–16465. 10.1073/pnas.0508239102

Chang, E. F., & Merzenich, M. M. (2003). Environmental Noise Retards Auditory Cortical Development. Science, 300(5618), 498–502. 10.1126/science.1082163

Churchland, M. M., Yu, B. M., Cunningham, J. P., Sugrue, L. P., Cohen, M. R., Corrado, G. S., Newsome, W. T., Clark, A. M., Hosseini, P., Scott, B. B., Bradley, D. C., Smith, M. A., Kohn, A., Movshon, J. A., Armstrong, K. M., Moore, T., Chang, S. W., Snyder, L. H., Lisberger, S. G.,… Shenoy, K. V. (2010). Stimulus onset quenches neural variability: A widespread cortical phenomenon. Nature Neuroscience, 13(3), 369–378. 10.1038/nn.2501

Collignon, O., Dormal, G., Albouy, G., Vandewalle, G., Voss, P., Phillips, C., & Lepore, F. (2013). Impact of blindness onset on the functional organization and the connectivity of the occipital cortex. Brain, 136(9), 2769–2783. 10.1093/brain/awt176

De Villers-Sidani, E., Chang, E. F., Bao, S., & Merzenich, M. M. (2007). Critical Period Window for Spectral Tuning Defined in the Primary Auditory Cortex (A1) in the Rat. The Journal of Neuroscience, 27(1), 180–189. 10.1523/JNEUROSCI.3227-06.2007

Dinstein, I., Heeger, D. J., & Behrmann, M. (2015). Neural variability: Friend or foe? Trends in Cognitive Sciences, 19(6), 322–328. 10.1016/j.tics.2015.04.005

Dosher, B. A., & Lu, Z. L. (1998). Perceptual learning reflects external noise filtering and internal noise reduction through channel reweighting. Proceedings of the National Academy of Sciences, 95(23), 13988–13993. 10.1073/pnas.95.23.13988

Glasberg, B. R., & Moore, B. C. J. (1990). Derivation of auditory filter shapes from notched-noise data. Hearing Research, 47, 103–138. 10.1016/0378-5955(90)90170-T

Goris, R. L. T., Movshon, J. A., & Simoncelli, E. P. (2014). Partitioning neuronal variability. Nature Neuroscience, 17(6), 858–865. 10.1038/nn.3711

Gougoux, F., Lepore, F., Lassonde, M., Voss, P., Zatorre, R. J., & Belin, P. (2004). Pitch discrimination in the early blind. Nature, 430(6997), 309–309. 10.1038/430309a

Huber, E., Chang, K., Alvarez, I., Hundle, A., Bridge, H., & Fine, I. (2019). Early blindness shapes cortical representations of auditory frequency within auditory cortex. Journal of Neuroscience, 39(26), 5143–5152. 10.1523/JNEUROSCI.2896-18.2019

Law, C.-T., & Gold, J. I. (2008). Neural correlates of perceptual learning in a sensory-motor, but not a sensory, cortical area. Nature Neuroscience, 11(4), 505–513. 10.1038/nn2070

Oxenham, A. J. (2012). Pitch perception. Journal of Neuroscience, 32(39), 13335–13338. 10.1523/JNEUROSCI.3815-12.2012

Park, W. J., & Fine, I. (2023). The perception of auditory motion in sighted and early blind individuals. Proceedings of the National Academy of Sciences, 120(49), e2310156120. 10.1073/pnas.2310156120

Patterson, R. D. (1976). Auditory filter shapes derived with noise stimuli. The Journal of the Acoustical Society of America, 59(3), 640–654. 10.1121/1.380914

Perrot, X., Ryvlin, P., Isnard, J., Guénot, M., Catenoix, H., Fischer, C., Mauguière, F., & Collet, L. (2006). Evidence for Corticofugal Modulation of Peripheral Auditory Activity in Humans. Cerebral Cortex, 16(7), 941–948. 10.1093/cercor/bhj035

Sabourin, C. J., Merrikhi, Y., & Lomber, S. G. (2022). Do blind people hear better? Trends in Cognitive Sciences, 26(11), 999–1012. 10.1016/j.tics.2022.08.016

Schütt, H. H., Harmeling, S., Macke, J. H., & Wichmann, F. A. (2016). Painfree and accurate Bayesian estimation of psychometric functions for (potentially) overdispersed data. Vision Research, 122, 105–123. 10.1016/J.VISRES.2016.02.002

Shadlen, M. N., & Newsome, W. T. (1994). Noise, neural codes and cortical organization. Current Opinion in Neurobiology, 4(4), 569–579. 10.1016/0959-4388(94)90059-0

Shera, C. A., Guinan, J. J., & Oxenham, A. J. (2002). Revised estimates of human cochlear tuning from otoacoustic and behavioral measurements. Proceedings of the National Academy of Sciences, 99(5), 3318–3323. 10.1073/pnas.032675099

Terreros, G., & Delano, P. H. (2015). Corticofugal modulation of peripheral auditory responses. Frontiers in Systems Neuroscience, 9. 10.3389/fnsys.2015.00134

Tomko, G. J., & Crapper, D. R. (1974). Neuronal variability: Non-stationary responses to identical visual stimuli. Brain Research, 79(3), 405–418. 10.1016/0006-8993(74)90438-7

Voss, P., & Zatorre, R. J. (2012). Occipital cortical thickness predicts performance on pitch and musical tasks in blind individuals. Cerebral Cortex, 22(11), 2455–2465. 10.1093/cercor/bhr311

Wan, C. Y., Wood, A. G., Reutens, D. C., & Wilson, S. J. (2010). Early but not late-blindness leads to enhanced auditory perception. Neuropsychologia, 48(1), 344–348. 10.1016/J.NEUROPSYCHOLOGIA.2009.08.016

Watkins, K. E., Shakespeare, T. J., O’Donoghue, M. C., Alexander, I., Ragge, N., Cowey, A., & Bridge, H. (2013). Early auditory processing in area V5/MT+ of the congenitally blind brain. The Journal of Neuroscience : The Official Journal of the Society for Neuroscience, 33(46), 18242–18246. 10.1523/JNEUROSCI.2546-13.2013

Zhang, L. I., Bao, S., & Merzenich, M. M. (2001). Persistent and specific influences of early acoustic environments on primary auditory cortex. Nature Neuroscience, 4(11), 1123–1130. 10.1038/nn745

